# A single-nuclei multiomics resource across four brain regions prioritises human neural cell types influencing brain-related traits

**DOI:** 10.64898/2026.09.24.754059

**Authors:** Ann C Babtie, Georgina E T Blake, Youth-GEMs Consortium, Nicholas E Clifton, Thomas Hawes, Inês Barroso, Jonathan Mill

**Author notes:** Joint lead authors.

## Abstract

Genetic studies have identified thousands of variants associated with brain-related traits. However, the majority of these map to non-coding regions and their causal roles and functional consequences are unclear. In this study, we profiled gene expression and chromatin accessibility in ∼140,000 individual nuclei from 40 post-mortem adult human brain samples from 11 donors spanning four brain regions (amygdala, hippocampus, hypothalamus and prefrontal cortex). Integrating these data with genome-wide association study statistics allowed us to prioritise specific neural cell populations relevant for complex traits. Concordant with epidemiological evidence, we prioritise similar neuronal cell populations for BMI, schizophrenia, bipolar disorder and age at menarche associated variants. Our paired multiomic data also provides support for putative enhancer-gene relationships relevant to Alzheimer’s disease. These data provide a valuable resource to help interpret trait-associated genetic variation and nominate effector transcripts and cellular pathways relevant to brain-related phenotypes.

## INTRODUCTION

The human brain is a highly complex organ with diverse specialised cell types and functionally distinct regions^1^. Advances in single-cell and spatial transcriptomic technologies have enabled detailed characterisation of neural cell types and their transcriptional states, with a predominant focus initially on cortical regions^2–6^. Increasingly, single-cell resolution data is now extending to other brain regions and incorporating epigenomic and proteomic data^1,3,7–14^. Epigenomic profiling to probe chromatin accessibility, chromosomal conformation, or DNA and histone modifications provides crucial information to better understand the regulatory elements and mechanisms that influence cell type-specific gene expression.

Using such data to improve functional genomic annotations across relevant tissues and cell types can aid the interpretation of trait-associated genetic variation^15–18^. Genome-wide association studies (GWASs) have identified thousands of common single nucleotide polymorphisms (SNPs) associated with brain-relevant traits, including psychiatric disorders, neurodegenerative diseases and non-neurological phenotypes influenced by brain function such as body mass index (BMI) ^19–21^. However, most trait-associated variants are located in non-coding regions and their causal roles and mechanisms of action are poorly understood^17,19,20^. Transcriptomic and epigenomic data across brain regions and cell types can help us prioritise causal variants and better understand their impacts in terms of cell types, pathways and genes affected^5,6,9,10,16–18,22^.

In this study, we generated cell type-specific transcriptomic and chromatin accessibility profiles across four brain regions (amygdala, hippocampus, hypothalamus and prefrontal cortex) using single-nuclei multiomic sequencing. We focused on regions implicated in homeostatic functions (e.g. appetite regulation by hypothalamic pathways) and those involved in reward circuitry and decision making (amygdala, hippocampus and prefrontal cortex) ^19,23^. These data contribute to ongoing efforts to characterise multiple brain regions and the functional and regional heterogeneity of neural cells and are available as a resource to the wider community. By integrating our dataset with trait-associated SNPs from GWAS, we prioritised cell types relevant for complex traits and identified shared enrichment across neuronal subtypes associated with BMI, age at menarche and psychiatric traits. Quantifying both chromatin accessibility and gene expression within the same nuclei allowed us to explore *cis*-regulatory mechanisms. We provide additional evidence for microglial-specific associations between putative regulatory elements harbouring Alzheimer’s disease (AD) risk variants and their candidate effector genes, highlighting the value of single-cell multiomic data in contributing to our mechanistic understanding of complex phenotypes.

## RESULTS

### Generation of single nuclei gene expression and chromatin accessibility profiles across four human brain regions

We jointly profiled gene expression and chromatin accessibility in individual nuclei isolated from four brain regions (amygdala, hippocampus, hypothalamus, prefrontal cortex) using single-nucleus multiome sequencing (see **Methods**). Brain tissue was obtained from 11 donors (aged 62-90 years, 6 female and 5 male) with minimal neuropathology and no known neurological disease, comprising 40 samples across four brain regions (n = 10 per region). All four regions were available for nine donors, while the remaining two donors collectively contributed four samples (Table S1). A total of 143,607 nuclei passed quality control including ambient RNA removal and filtering based on both RNA and ATAC data quality metrics (see Methods, Fig. S1). Following integration of RNA data from all samples using Harmony^24^ nuclei were clustered based on gene expression into eight groups corresponding to major neural cell types (Fig. 1A, Fig. S2). Cell types were assigned to each cluster based on the expression of established neural marker genes (Table S2) ^1,22,25–28^ identifying astrocyte, ependymal, excitatory neuron, inhibitory neuron, microglia, oligodendrocyte, oligodendrocyte precursor (OPC) and vascular cells (Fig. 1B-C, Fig. S3). These manual cell type assignments were subsequently validated by mapping nuclei to the Allen Institute’s whole human brain reference taxonomy (Fig. S4) ^1^.

**Figure 1:**
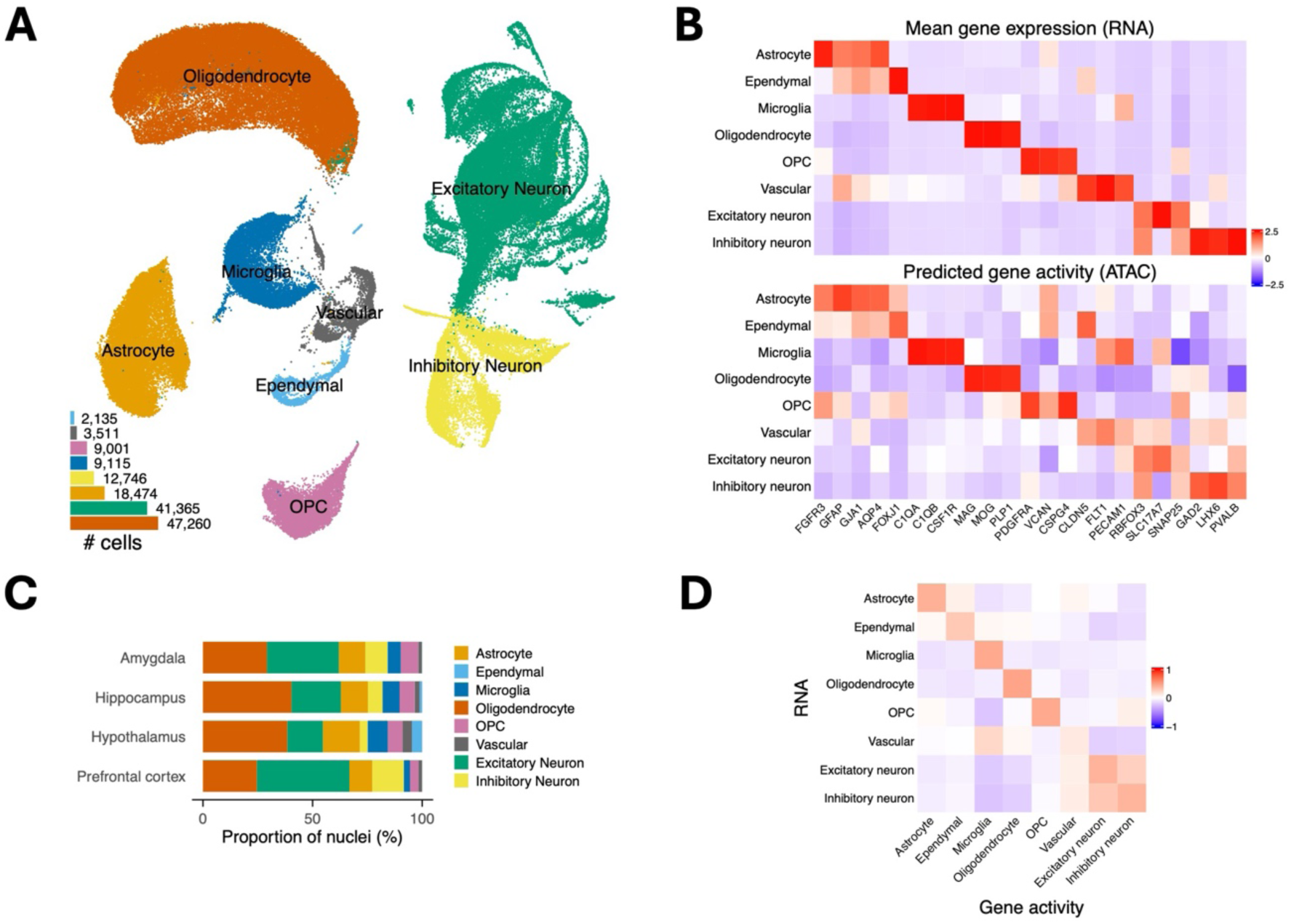
Overview of the single-nuclei multiome dataset across four human brain regions. (A) UMAP visualisation of snRNA-seq data coloured by annotated cell type (*n* = 143,607 nuclei); inset bar graph shows total number of nuclei per cell type. (B) Heatmaps showing mean gene expression level (top) and mean predicted gene activity derived from chromatin accessibility data (bottom) for selected marker genes across cell types. Colour indicates z-scaled mean expression or predicted activity per gene. (C) Percentage of total nuclei sampled from each brain region annotated to each cell type. (D) Pearson correlation between mean gene expression and predicted gene activity (derived from chromatin accessibility data) for each cell type.

Chromatin accessibility quantified within the same nuclei identified cell-type specific regions of open chromatin. We used the RNA-based cell type annotations and MACS3^29^ to create a consensus peak set comprising 670,427 peaks. 59 % of these peaks were unique to a single major cell type and 33 % of peaks were restricted to neurons (Fig. S5). Cell type-specific chromatin accessibility across genic regions correlated with the corresponding cell type-specific gene expression profiles (Fig. 1D). For matched cell types, the correlations across modalities were higher (range of 0.11-0.47, median = 0.40) than those observed between non-matched cell types (range –0.25-0.29, median = - 0.06). Joint analysis of both data modalities using a weighted-nearest neighbour approach^30^ indicated that RNA features were more informative than ATAC features in defining cell state for most nuclei (87.6 %; Fig. S6). We therefore retained the RNA-based cell type annotations for all downstream analyses.

### Gene expression varies across neuronal sub types and brain regions

Neural cells show high transcriptional heterogeneity across cell types and brain regions reflecting their diverse and specialised functions^1^. To quantify the sources of this variation we used variance partitioning to assess the contribution of technical and biological factors to gene expression variability^31,32^. Cell type was the dominant factor associated with variation in gene expression (Fig. 2A, Fig. S7), consistent with other human brain single cell studies of the human brain^1,5^. Across all genes, a mean of 54.6 % of total expression variance was attributed to cell type, 3.9 % to individual and 38.6 % to unexplained residual variation, with less than 1 % attributed to the other tested factors (Table S3). As expected, known cell type marker genes displayed high cell type-associated variance and sex chromosome genes showed the highest sex-associated expression variance (Fig. S8).

**Figure 2:**
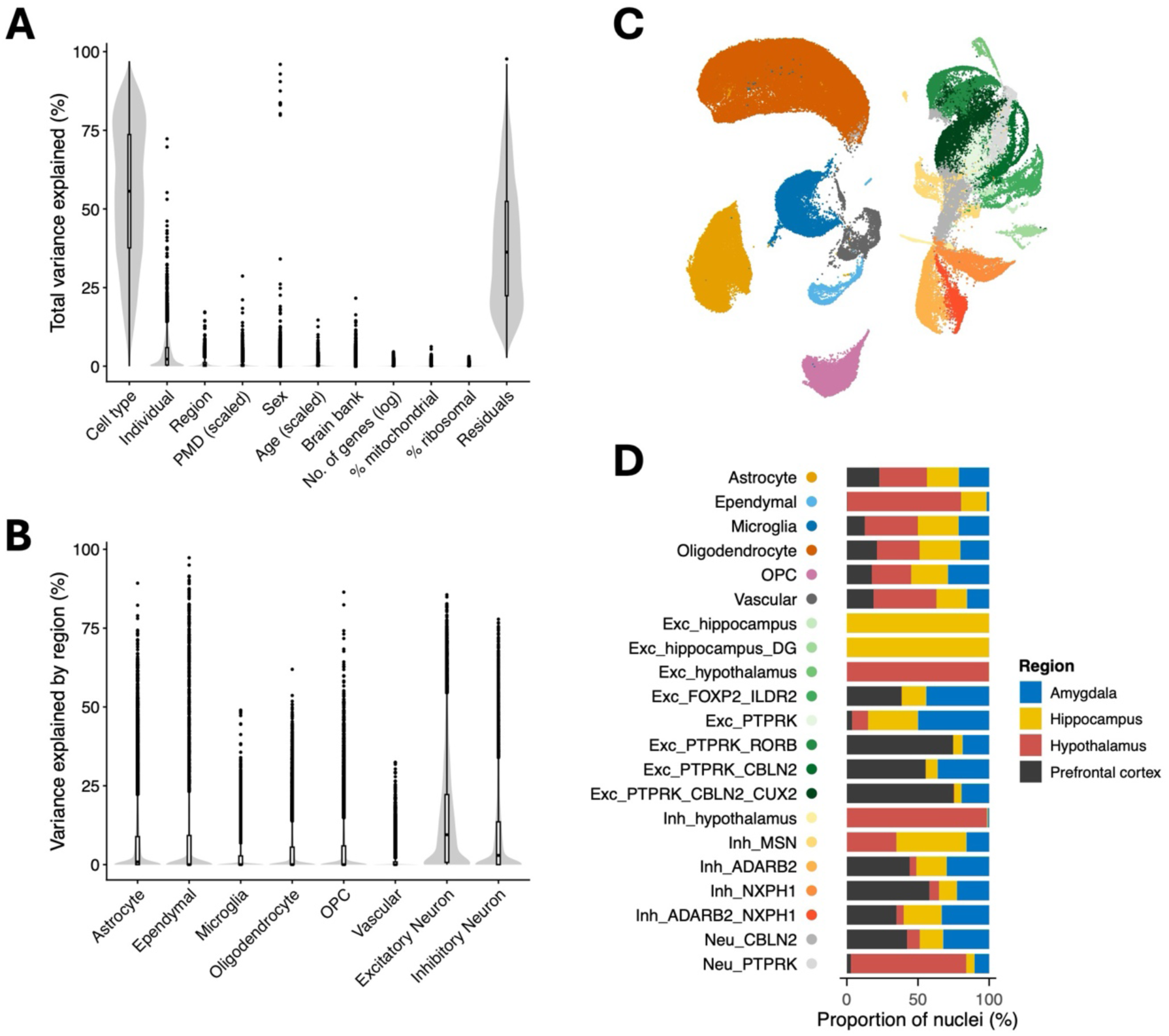
Gene expression variability across cell types and brain regions. (A) Proportion of gene expression variance (*n* = 11,666 genes) explained by selected covariates. Box plots within violin plot distributions indicate the median and interquartile range (IQR), with whiskers extending to the largest and smallest values within 1.5xIQR from the box ends; outlier points beyond these values are indicated by dots. (B) Proportion of variance explained by brain region in gene expression data separated by cell type. (C) UMAP visualisation of snRNA-seq data coloured by major cell types (non-neuronal) and finer neuronal subtype classifications (D) Proportion of nuclei assigned to each cell type and neuronal subtype across the four sampled brain regions. Filled coloured circles next to cell type names in Fig 2D indicate the colours used in Fig 2C for cell (sub)types.

Brain region accounted for only a small proportion of overall transcriptomic variation, but when cell types were analysed separately, region explained a mean of 14.3 % and 9.2 % of total gene expression variation in excitatory and inhibitory neurons, respectively (Fig. 2B, Table S4). To further dissect neuronal diversity, we re-clustered neuronal nuclei at increased resolution to identify subgroups with distinct gene expression profiles (Fig. 2C, Fig. S9). These subgroups were annotated using reference-based labels of their constituent nuclei, expression patterns of genes used for annotation of the reference human brain taxonomy^1^, and their distributions across sampled brain regions (Fig. 2D, Fig. S10). While some neuronal subtypes were detected across all four regions, others exhibited pronounced regional diversity with four neuron subtypes (Exc-hippocampus, Exc-hippocampus-DG (dentate gyrus), Exc-hypothalamus, and Inh-hypothalamus) comprised of nuclei originating almost entirely (> 98 %) from a single region (hippocampus or hypothalamus) (Fig. S11).

### Mapping trait associations across neural cell-types

We integrated cell type specific gene expression and chromatin accessibility profiles with GWAS summary statistics for 27 neurodegenerative, psychiatric, metabolic, reproductive and anthropometric traits (Table S5) to prioritise relevant neural cell types. Using MAGMA gene-property analysis^33^, we tested whether genes associated with each trait were preferentially expressed in specific cell types (Fig. 3A, Fig. S12, Tables S6-7). As expected, microglia showed a significant association with Alzheimer’s Disease (AD) (β = 0.31, FDR = 4.4x10^-9^, in major cell type analysis) consistent with their established role in AD pathogenesis^34,35^.

**Figure 3:**
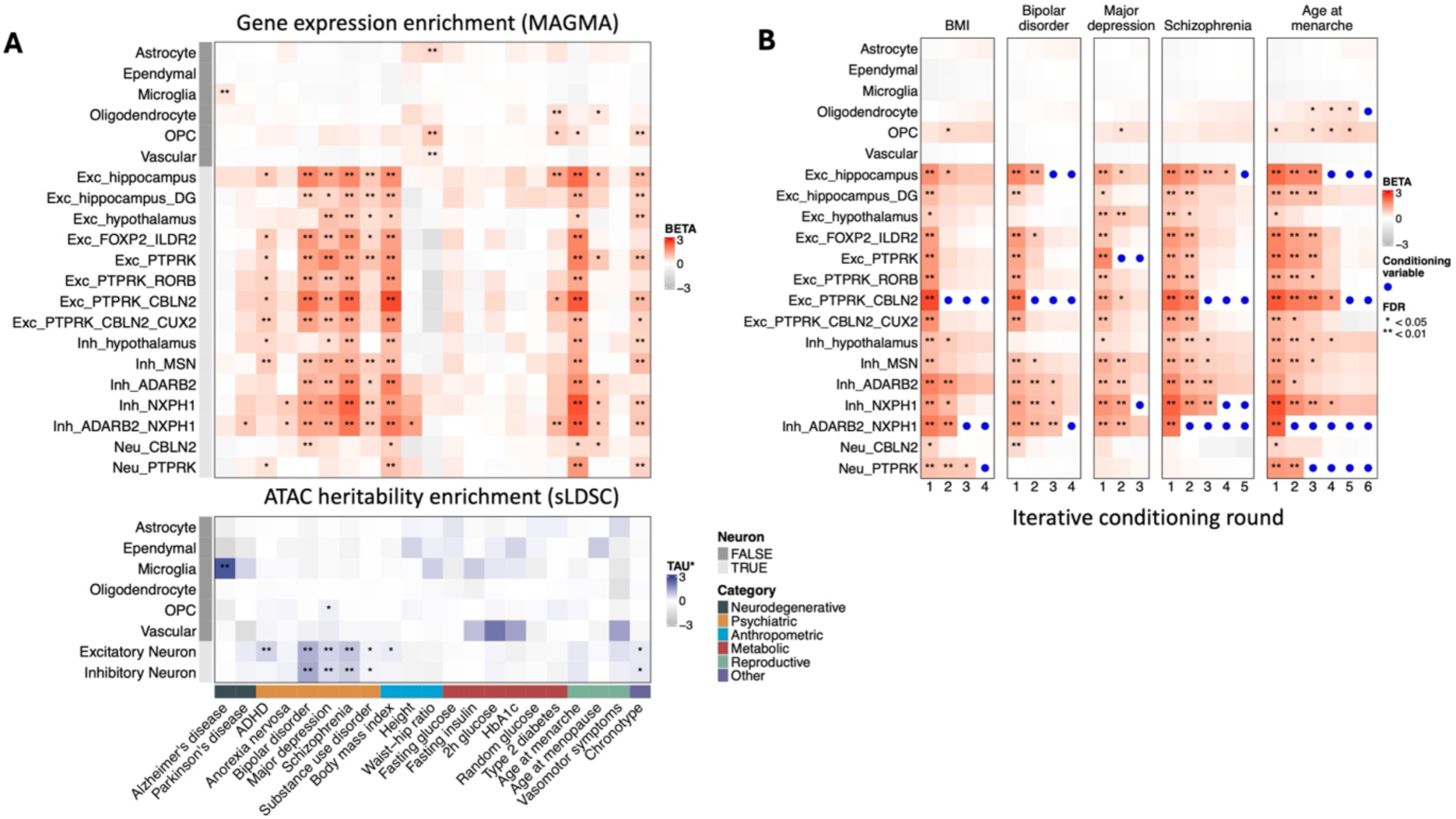
Associations between cell type specific gene expression, chromatin accessibility, and brain-related traits. (A) *Upper panel*: MAGMA gene property analyses testing association between traits and increased gene expression specificity across cell types. Heatmap shows effect sizes (*beta*) for one-sided enrichment tests; * and ** indicate FDR-corrected *p*-values < 0.05 and 0.01, respectively. *Lower panel*: Stratified LD score regression (sLDSC) heritability enrichments within cell type specific open chromatin regions. Heatmap shows standardised effect sizes (*tau*\*); * and ** indicate FDR-corrected one-sided *p*-values < 0.05 and 0.01, respectively, calculated from LDSC coeffiicient *z*-scores. (B) Conditional MAGMA gene property analyses. For each trait, the first column reproduces the unconditioned results shown in (A). Subsequent columns show the results from iterative rounds of conditioned analyses where the most significant cell type (lowest FDR) from the prior round is added to the conditioning set. Filled blue circles indicate the cell types included as conditioning variables in each round. Multiple testing correction was performed across cell types within each trait for both MAGMA and sLDSC analyses.

Multiple neuronal subtypes were significantly associated with five psychiatric traits (ADHD, bipolar disorder, major depression, schizophrenia, substance use disorder), in addition to BMI, age at menarche and chronotype with > 8 significant neuronal subtypes identified for each trait; Fig. 3A, Tables S6-7). Conditional MAGMA analyses were used to evaluate whether these associations reflect a shared neuronal expression specificity signature or were driven by independent neuron subtype-specific transcriptional profiles. For the five traits showing the broadest neuronal associations (bipolar disorder, major depression, schizophrenia, BMI, and age at menarche), we conducted an iterative conditional analysis in which the most significant cell type from each round was added to the set of conditioning variables for the next round (Fig. 3B). These analyses indicated that the observed trait associations were partly driven by expression signatures shared across neuronal subtypes, but also included independent contributions from subtype-specific transcriptional profiles.

For BMI, the strongest association in the unconditioned analysis was for the excitatory neuron subtype Exc-PTPRK-CBLN2 (β = 2.62, FDR = 1.7x10^-8^). Conditioning on this subtype substantially attenuated associations across most other excitatory neuron subtypes and several inhibitory neuron subtypes (Fig. 3B) consistent with a shared expression-specificity profile. Supporting this interpretation, these subtypes showed relatively high correlations with the expression specificity profiles of Exc-PTPRK-CBLN2 (Fig. S13). However, the association between Exc-PTPRK-CBLN2 cells and BMI remained significant after conditioning on other cell types independently (Fig. S14), indicating an additional subtype-specific component to the association. Subtype-specific contributions were also observed for Inh-ADARB2-NXPH1 and Neu-PTPRK (Fig. 3B), with Exc-PTPRK-CBLN2 and Inh-ADARB2-NXPH1 showing these independent contributions consistently across bipolar disorder, schizophrenia, age at menarche and BMI, highlighting overlap in the neuron subpopulations implicated across these traits.

Type 2 diabetes (T2D) was the only metabolic trait tested that showed significant associations with neural cell types. T2D-associated genetic variants have previously been grouped into mechanistic clusters based on their broader phenotypic associations^36^. We therefore used Expression Weighted Celltype Enrichment (EWCE)^37^ to assess whether specific mechanistic clusters accounted for the observed neural cell type associations. Inhibitory neurons and oligodendrocyte precursors showed significantly enriched expression of genes associated with the obesity-associated T2D mechanistic cluster (Fig. S15), with lower neural cell type enrichments observed across other mechanistic clusters. These findings suggest that the observed T2D associations in Fig. 3A are more strongly related to obesity-associated mechanisms than glycaemic regulation per se.

### Heritability enrichment within cell type-specific regulatory landscapes

To complement the MAGMA analyses we used stratified LD score regression (sLDSC)^38–40^ to test whether SNP-heritability, i.e. the proportion of phenotypic variation attributed to trait-associated common variants, is enriched in cell type-specific genomic annotations derived from gene expression and chromatin accessibility data (see Methods). Heritability enrichments within cell type-specific accessible chromatin annotations broadly recapitulated the strongest cell type-trait associations identified by MAGMA, including enrichment in microglia for AD, and in excitatory and/or inhibitory neurons for ADHD, bipolar disorder, depression, schizophrenia, substance use disorder, BMI and chronotype (Fig. 3A, Fig. S16).

### Linking regulatory elements to target genes implicated in Alzheimer’s disease

We leveraged paired RNA and ATAC data to nominate candidate effector genes for trait-associated variants by modelling covariation between chromatin accessibility and gene expression. We used pgBoost to generate consensus cell type-specific scores for candidate SNP-gene links by integrating predictions from three peak-to-gene or peak-to-peak linking methods (SCENT, Signac and Cicero) with distance-based metrics^41–44^. PgBoost models were trained using fine-mapped expression quantitative trait loci (eQTL) data from GTEx brain tissues^45^ and used to score candidate links across six cell types (astrocytes, oligodendrocytes, oligodendrocyte precursors, microglia, and excitatory and inhibitory neurons). We evaluated pgBoost predicted SNP-gene links across different validation datasets expected to be enriched in true variant-gene pairs; these included human brain cell type-specific eQTL and colocalisation data^22,46^ and SNP-gene pairs prioritised for brain-related traits (BMI, AD, schizophrenia) by the Open Targets Locus-to-Gene algorithm^47,48^ (Table S8). Across all validation datasets these SNP-gene pairs were significantly enriched among high-scoring pgBoost predictions (Fig. S17, Table S8), demonstrating that our single nucleus multiomic data captures signals informative for linking trait-associated genetic variants to putative effector genes.

To illustrate the utility of these data for prioritising variant-gene links, we examined AD-associated risk loci in greater detail (Fig. 4 and Fig. S18). Fourteen links within the top 5 % of pgBoost scores included an AD credible set lead variant reported by Open Targets. For each, the top scoring candidate target gene matched the gene prioritised by the Open Targets Locus to Gene algorithm^47^. Eight of these variants overlap microglial-specific accessible chromatin peaks, with their predicted target genes also preferentially expressed in microglia, consistent with cell type-specific enhancer-gene regulatory relationships. These links were further supported by independent evidence implicating the corresponding genes in AD, as detailed below and in Fig. S18.

**Figure 4:**
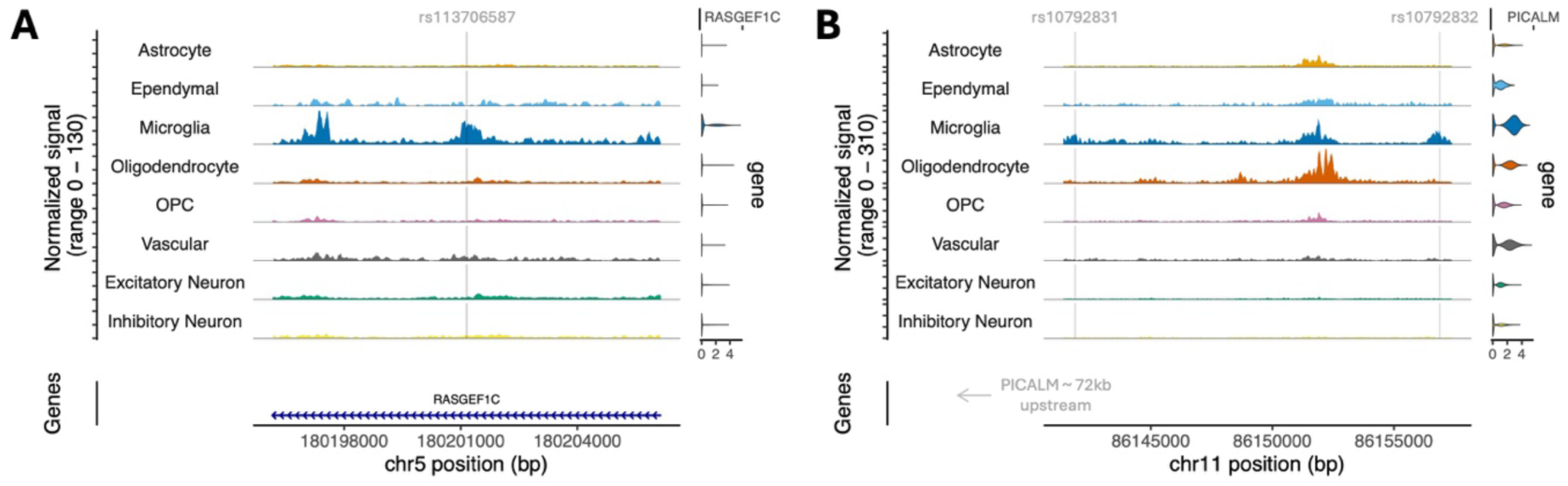
Linking Alzheimer’s disease risk variants to candidate target genes. (A-B) Coverage plots showing normalised chromatin accessibility signal across major cell types at selected genomic regions. Tracks are centered on regions surrounding genetic variants prioritised by pgBoost, and violin plots (right) show cell-type specific gene expression profiles of the corresponding target genes. In each case the variants represent the lead SNP from an Alzheimer’s disease GWAS credible set identified by Open Targets and the same genes are prioritised by Open Targets Locus-to-Gene predictions. Variants are indicated by the vertical grey lines: rs113706587 (chr5:180201150, hg38), rs10792831 (chr11:86141937) and rs10792832 (chr11: 86156833).

Variant rs113706587 (Fig. 4A) has been associated with early amyloid deposition using positron emission tomography image analysis^49^, and *RASGEF1C* was previously identified as an AD-relevant gene in transcriptomic, DNA methylation and genetic analyses^5,50,51^. A single nuclei eQTL study found colocalisation of microglial *RASGEF1C* eQTL and AD-risk loci with fine-mapping of the *RASGEF1C* eQTL locus prioritising rs113706587 as a likely causal variant^46^. Consistent with this, a deep learning model predicted that rs113706587 disrupts a microglial accessible chromatin peak, and CRISPR-based targeting of this putative regulatory element in induced pluripotent stem cell (iPSC)-derived microglia reduced *RASGEF1C* expression^52^.

Two AD-associated variants (rs10792831 and rs10792832) were linked to *PICALM* in our prioritised variant-gene links (Fig. 4B). Both variants have been predicted by deep learning models to influence chromatin accessibility^52,53^. Epigenomic annotations indicate these variants fall in putative microglial-specific enhancers that have been linked to the *PICALM* promoter region via proximity ligation-based assays^54^. AD risk variant rs10792832 colocalises with a microglial *PICALM* eQTL^46,55^ and at heterozygous loci is associated with an allelic imbalance of accessible chromatin^56^. Recently, evidence for a causal link between the AD risk allele of variant rs10792832 and reduced *PICALM* expression via the disruption of transcription factor *PU.1* binding was established in human iPSCs, resulting in aberrant lipid droplet accumulation and phagocytic function in microglia^56^. Together, these examples (Fig. 4 and Fig. S18) illustrate how cell type-resolved multiomic data can prioritise candidate effector genes and provide mechanistic insight into non-coding trait-associated variants.

## DISCUSSION

In this study, we generated a single nuclei multiomic atlas of gene expression and chromatin accessibility across four regions of the human brain. Integrating these data with genetic association studies highlighted distinct neural cell populations relevant to different brain-related phenotypes, including shared neuronal subtypes implicated across bipolar disorder, schizophrenia, age at menarche and BMI, and microglia-specific enhancer-gene relationships at AD risk loci.

This data resource complements ongoing efforts^1,7,9–11,22,57^ to extend transcriptional and epigenomic profiling of human neural cells beyond the cortex to include the amygdala, hippocampus and hypothalamus, brain regions that have been comparatively underrepresented in existing atlases. Joint measurement of gene expression and chromatin accessibility within individual nuclei further enables the investigation of cell-type-specific regulatory mechanisms and their relationship to human disease. Our data are available as a resource for the broader research community, facilitating the exploration of cell type-specific molecular features and the interpretation of genetic associations.

Although cell type represented the dominant source of transcriptomic variation across the dataset, substantial regional heterogeneity was evident within neuronal populations. The high transcriptional diversity observed amongst neurons is consistent with previous studies describing extensive heterogeneity in neuron gene expression regulation and regional specialisation^1,5,7,9,58^. In contrast to glial populations, which are characterised by comparatively modest regional differences, excitatory and inhibitory neurons showed marked region-associated transcriptional variation, with several neuronal subtypes specific to either the hippocampus or hypothalamus.

Integration with GWAS studies prioritised neural cell populations relevant to brain-related phenotypes. Consistent with previous studies, we observed strong enrichment of AD risk within microglia^7,16,46,59,60^, and broad excitatory and inhibitory neuronal associations with BMI^9^, psychiatric traits^16,18,59,60^, age at menarche, and chronotype. These results suggest that the genetic architecture of these traits is mediated through molecular programmes shared across multiple neuronal subtypes, rather than being restricted to a small number of specialised cell populations. Notably, conditional analyses also identified evidence for subtype-specific contributions, indicating that both shared and unique neuronal transcriptional features contribute to the observed enrichments. In line with established differences in power^18,61^, complementary ATAC- and RNA-based heritability enrichments using sLDSC were consistent for the strongest cell type-trait associations (Fig. 3A, Fig. S16).

Specific neuron subpopulations implicated for schizophrenia associations (excitatory subtypes Exc-hippocampus and Exc-PTPRK-CBLN2 and inhibitory subtypes Inh-NXPH1 and Inh-ADARB2-NXPH; Fig. 3B) are analogous to subtypes prioritised by published conditional analyses^16,18^ (namely amygdala excitatory, hippocampal CA1-3 and deep-layer intratelencephalic excitatory subtypes, and CGE interneuron inhibitory subtypes; Fig. S10). Conditional analyses revealed shared neuron subtypes prioritised for BMI, bipolar disorder, schizophrenia and age at menarche, and to a lesser extent multiple depression (Fig. 3B). Common cell type enrichment profiles suggest shared cellular processes may influence these diverse phenotypes and could contribute to the epidemiologically observed comorbidities amongst these traits^62–67^.

For T2D, a subset of obesity-related T2D genetic variants appear to drive the neuron subtype and oligodendrocyte lineage associations (Fig. S15). These findings are consistent with previous ATAC-based analyses that also found neural cell type associations with obesity-related mechanistic subsets of T2D genetic risk variants^36^. However, their more widespread associations extending to metabolic syndrome related variants are not fully replicated by our RNA-based analyses which may be a limitation of annotating genes to variant clusters based on proximity. Nevertheless, these results distinguish distinct subsets of T2D variants likely influencing phenotype via neural cell processes and contributing to the observed phenotypic heterogeneity of T2D.

Variant-target gene links predicted from our paired cell type specific accessibility and expression profiles were enriched for variant-gene pairs in validation datasets (Fig. S17). While demonstrating that single nuclei multiomic data are informative, particularly when integrated alongside distance metrics, benchmarking studies show predictions often lack concordance across different statistical implementations^43,44^. As such these methods are best viewed as mechanisms to generate hypotheses about putative target genes or contribute additional evidence and cell type context in support of proposed variant-gene links, rather than expecting them to reliability nominate biologically relevant relationships independently. We selected top-scoring variant-gene links for AD-risk variants annotated as credible set lead SNPs (Fig. 4) ^48,68^; these examples show microglial specific variant accessibility and corresponding target gene expression, providing cell type context and further support for these proposed regulatory element-gene relationships.

Our results demonstrate the value of this dataset, while also highlighting opportunities for future expansion. First, increasing the number of donors and incorporating genotype data would enable cell type specific molecular QTL analyses, providing more direct insight into the regulatory consequences of genetic variation and supporting causal inference approaches^17,22,26,46,69^. Although profiling neurologically unaffected donors provides a valuable reference for molecular variation in the absence of disease-related changes^22^ larger and more diverse cohorts would enable case-control analyses and more systematic exploration of the influence of sex, developmental stage and/or ancestry, the latter particularly relevant in the context of current efforts to address inequalities in genomic analyses^70^. Incorporating additional data modalities would improve characterisation of genomic regulatory elements and their downstream effectors. Long-read sequencing of transcript isoforms may be particularly informative for neurological phenotypes given the extensive alternative splicing observed in the brain^71,72^ while emerging single-nucleus approaches enable characterisation of DNA and histone modifications^8,12,73–75^.

In conclusion, we present a single nuclei data resource integrating transcriptomic and chromatin accessibility profiles across multiple brain regions. We illustrate the value of these data by highlighting neural cell types and putative regulatory relationships relevant for selected brain-related traits. These data provide a reference for interpreting trait-associated genetic variation in in a cell-type- and region-specific context and contributes to ongoing efforts to define the molecular mechanisms underlying complex traits.

## METHODS

### Sample details

Post-mortem tissue from 11 adult donors (aged 62-90 years, 6 females, 5 males) with no neuropathology or diagnosed neurodegenerative disease was obtained from the MRC UK Brain Bank Network (the University of Oxford Brain Bank (OBB) and the South West Dementia Brain Bank (SWDBB)). Tissue was provided by the OBB and SWDBB under their respective NHS REC generic approval. A total of 40 tissue samples were used, 10 from each of four brain regions (amygdala, hippocampus, hypothalamus and prefrontal cortex). Table S1 provides an overview of the samples.

### Single nuclei multiome assays

Single nuclei suspensions were obtained from ∼50mg of post-mortem brain tissue from each brain region. Briefly, tissue was homogenised in a pre-chilled dounce homogeniser prior to nuclei isolation via ultracentrifugation over an 8M sucrose cushion. Isolated nuclei were then washed and resuspended in PBS and 2% BSA (a full protocol is provided on protocols.io at https://dx.doi.org/10.17504/protocols.io.eq2lym7prlx9/v1). Ribolock RNase-inhibitor (Thermo Scientific, EO0382) was added to the lysis and resuspension buffers (0.4 U/ul). Nuclei suspensions were assessed for the presence of debris and manually counted on a haemocytometer. Transposition, GEM generation and library preparation was carried out as per the 10x Genomics Chromium Next GEM Single Cell Multiome ATAC + Gene Expression user guide (CG000338 Rev F). Targeted recovery of 4000 nuclei per sample was used. cDNA and final library quantification, quality control and fragment size determination was performed using the D5000 high sensitivity ScreenTape assay and reagents (Agilent technologies). Gene expression and ATAC-seq libraries were pooled and sequenced on an Illumina NovaSeq6000 machine.

### Raw data processing

10x Genomics Cell Ranger ARC (v2.0.2) was used to align sequencing reads to reference genome GRCh38 (with GENCODE v32 reference annotation) and generate feature barcode count matrices. CellBender (v0.3.0)^76^ was used to remove ambient RNA and generate filtered feature count matrices (excluding ATAC peak features from analysis); 0.39-1.57 % of RNA counts were removed from non-empty droplets per sample.

### Quality control of single nuclei multiome data

R (v4.3.3) and R packages Seurat (v5.1.0)^77^ and Signac (v1.13.0)^42^ were used for downstream analysis. We excluded RNA features detected in fewer than 5 nuclei, and removed data from nuclei barcodes meeting the following criteria: < 200 RNA counts, < 200 ATAC counts, > 25 % RNA counts mapping to mitochondrial genes, nucleosome signal > 3 (ratio of mono-nucleosomal to nucleosome-free fragments), or those identified as doublets using scDblFinder^78^.

### Clustering and cell type annotation

Cell type annotation was based on the gene expression data. We first used Seurat to log normalise and calculate principal components (PCs) based on 4,000 highly variable genes before integrating samples using Harmony^24^. The top 30 Harmony-corrected PCs were used to cluster nuclei using the Leiden algorithm and generate a uniform manifold approximation and projection (UMAP) embedding. We labelled clusters corresponding to the following neural cell types based on expression of previously reported marker genes: astrocytes (*AQP4*, *FGFR3*), ependymal (*FOXJ1*), excitatory neurons (*RBFOX3*, *SLC17A7*, *SYT1*), inhibitory neurons (*GAD1*, *GAD2*), microglia (*C1QB*, *CSF1R*), oligodendrocytes (*MOBP*, *PLP1*), oligodendrocyte precursors (*PDGFRA*, *VCAN*), and vascular (*FLT1I*, *PECAM1*); the full list of marker genes is given in Table S2 ^1,22,25,27,28,79^. Manual annotations were verified by comparison to cell type labels generated by the Allen Brain Institute’s MapMyCells resource (RRID:SCR_024672) (Fig. S4). Neuronal nuclei were selected and Leiden clustering repeated to identify transcriptionally similar subgroups. Neuronal subgroups were assigned labels based on expression patterns of genes used for auto-annotations in Siletti *et al* ^1^ and comparison to the MapMyCells annotations (Fig. S9-10); labels indicate either regional or gene expression specificity of that group of nuclei (Fig. 2D, Fig. S9).

### ATAC data processing and peak calling

Sample specific peak sets called by CellRanger ARC were combined to create a common peak set by merging intersecting peaks and retaining peaks on standard chromosomes with widths between 50-5000 bp. This common feature set was used to merge ATAC data from all samples and create ATAC fragment sets for each major cell type cluster. Cell type specific peaks were called using MACS3^29^ (paired-end mode, q-value threshold = 0.05) and merged to create a consensus peak set. These MACS3 consensus peaks were quantified in the merged ATAC data before latent semantic indexing (LSI) to normalise peaks and reduce dimensions using the top 90 % most frequently observed peaks^42^. Harmony was used to correct the 2-50^th^ LSI components for sample batch effects (the first component was excluded as it correlated with sequencing depth). Weighted-nearest neighbour (WNN) analysis^30^ was used to create a joint neighbour graph representing both RNA and ATAC data based on the top 30 Harmony-corrected PCA and LSI components respectively. Gene activity estimates were calculated from ATAC-seq data based on counts per nuclei within gene bodies and promoter regions (+2 kb upstream of TSS).

### Cell type specificity scores

Pseudobulk RNA and ATAC count matrices were generated by summing raw counts from all nuclei within each annotated cell type cluster, and normalising to counts per million (CPM) within each cell type. We quantified cell type specificity using Expression Proportion (EP) – the proportion of a gene’s total (normalised) expression across all cell types that is found within a given cell type^18,61^; analogous peak specificity scores were calculated from pseudobulk ATAC data. For enrichment analyses requiring feature sets for each cell type, we used the top 10% of specificity scores per cell type to select gene or peak sets representative of that cell type, in line with previous recommendations^18,61^.

### GWAS summary statistics

Genome Wide Association Study (GWAS) summary statistics were downloaded for 26 traits from sources detailed in Table S5^36,68,80–97^. These span anthropometric, metabolic, neurodegenerative, reproductive, psychiatric and reproductive traits chosen to include both brain-related and non-brain related phenotypes. We filtered data to exclude variants with INFO score < 0.6 (where reported) and minor allele frequency < 0.01. MungeSumStats^98^ was used to standardise summary statistics, lift over genomic coordinates to GRCh37 where required (with UCSC hg38ToHg19 chain file), and exclude indels and non-autosomal variants.

### MAGMA cell type heritability enrichment

MAGMA (v1.10)^33^ gene-set and gene property association analyses were used to prioritise cell types relevant for each GWAS trait. First, gene level association statistics were calculated from the *P*-values of SNPs nearby each gene using MAGMA’s SNP-wise mean model. We restricted our analysis to protein-coding genes, used a window of 35 kb upstream to 10 kb downstream of each gene to include regulatory regions, excluded the extended major histocompatibility complex (xMHC) region (chr6: 25-34 Mb)^99^, and used the European 1000 Genomes Project Phase 3 reference panel^100^ to control for linkage disequilibrium (LD). Competitive gene-set analyses tested whether the mean association with a given phenotype was greater for genes within a gene set versus those outside. One-sided gene property analyses used continuous gene specificity scores to test if association with the phenotype was greater for specifically expressed genes. Both types of analyses controlled for gene size, gene density, and variation in sample sizes between SNPs, and conditioned analyses on the set of (or continuous CPM expression values for) brain-expressed genes (those expressed in any cell type in our data). *P-*values were adjusted for multiple comparisons across cell types within a trait using (Benjamini and Hochberg) FDR correction. Figures were created using the R package ComplexHeatmap (v2.18.0)^101^.

For cases where significant enrichments were observed across multiple neuronal subtypes we used conditional analyses to evaluate whether these associations are independent or driven by gene specificity profiles common to several subtypes. Two types of conditional gene property analysis were used for each trait: 1) association was tested in each cell type conditioned on gene specificities in each other cell type in turn; 2) an iterative approach where in each successive round, the most significant cell type (lowest FDR) from the previous round of analysis was added to a set of cell types to condition on for the next iteration, starting with the original gene property analysis (with no conditioning on cell types) and continuing until no significant associations remained.

### sLDSC cell type heritability enrichment

Stratified LD score regression (sLDSC) (v1.0.1)^38–40^ was used to test whether heritability for a GWAS trait is enriched in regions surrounding the top 10 % of specifically expressed genes or within the top 10 % accessible chromatin peaks in each cell type. We used LDSC’s *munge_sumstats.py* to further format GWAS summary statistics and restrict to HapMap3^102^ SNPs. Gene expression analysis was restricted to protein-coding genes excluding the xMHC region and used a window of +/-100 kb around each gene to define genomic annotations. Chromatin accessibility analysis used the top 10 % specific peak regions with no additional window. We used the 1000 Genomes Phase 3 LD reference panel and conditioned on the 53 genomic annotations in LDSC’s ‘baseline model’ ^40^ as well as the set of all brain-expressed genes (+/-100 kb) or the consensus set of MACS3 called peaks for the RNA/ATAC analyses respectively. We calculated the standardised effect size (*tau\**) – as recommended^103^ – to calculate enrichment *P*-values, and FDR correction across cell types within a trait.

### Expression weighted cell type enrichment (EWCE)

EWCE^37^ analysis tests if a gene list of interest shows enriched expression in particular cell types using single cell expression data. We excluded genes with < 5 total counts in our data, then used EWCE to remove non-variably expressed genes and calculate cell type gene specificity scores (based on mean expression of a gene across nuclei within a given cell type). We tested for enriched expression of selected gene sets using 10,000 bootstrap samples and the set of variable brain-expressed genes as background, controlling for GC content and transcript length. Our gene sets of interest were obtained from Suzuki *et al*. ^36^ where type 2 diabetes (T2D) GWAS variants were partitioned into 8 mechanistic clusters based on associations with other cardiometabolic traits. For each T2D mechanistic cluster we obtained a list of genes where at least one cluster-associated variant overlapped a defined genomic window (-35 kb/+10 kb or -10 kb/+2 kb) around the gene body (note these gene sets are overlapping). FDR corrected enrichment *P*-values were calculated across all cell types and gene sets tested.

### Variance partition (VP) analysis

R package dreamlet (v1.6.0)^31,32^ was used to evaluate the proportion of gene expression variance attributable to selected biological and technical covariates. We used the default workflow including aggregating data to the pseudobulk-level by summing raw counts across nuclei for each major cell type and sample combination, filtering genes by expression level (at least 5 counts in 40 % of pseudobulked samples), and voom-style normalisation. We defined the following regression formula for use in its linear mixed model framework: Gene expression ∼ (1|cell type) + (1|individual) + (1|region) + (1|BrainBank) + (1|Sex) + scale(Age) + scale(PMD) + log(nFeature_RNA) + percent.mt + percent.ribosomal

The categorical variables cell type, individual (donor), brain region, brain bank and sex are modelled as random effects, with the remaining variables as fixed effects; age and post-mortem delay (PMD) are scaled; and the mean across cells within a given cell type and sample was used for the cell-level covariates of number of genes detected (nFeature_RNA), and percentage of mitochondrial and ribosomal transcripts (percent.mt and percent.ribosomal). For cell type analysis, the initial term (cell type) was removed.

### Feature linking

Candidate links between single nucleotide variant (SNV) and target genes were prioritised using pgBoost^44^. This method generates consensus SNV-gene linking scores by combining variant-gene scores derived from single nuclei multiome data and genomic distance metrics using a gradient boosting framework. We generated cell-type specific peak-gene scores using Signac^42^ and SCENT^43^ and peak-peak scores using Cicero^41^; we restricted these analyses to six major cell types (astrocytes, microglia, oligodendrocytes, oligodendrocyte precursors, excitatory neurons, inhibitory neurons) and features (peaks or genes) that were detected in a minimum of 2 % or 5 % of cells in a given cell type (for Cicero/Signac and SCENT respectively). Following recommendations in ref. ^44^, candidate peak-gene links were restricted to *cis* peak-gene pairs with peak-transcription start site distances between 1-500 kb to prioritise enhancer-gene links, and we derived variant-gene scores from peak-gene scores based on 1000 Genomes project variants (minor allele count >= 5) overlapping each peak.

PgBoost scores were generated using a gradient boosting decision tree classification algorithm^104^ trained on fine-mapped GTEx eQTL data from 13 brain tissues using a leave-one-chromosome-out approach^44,45^. We restricted our analysis to protein-coding genes and defined positive and negative SNV-gene links as those with maximum posterior inclusion probabilities of > 0.2 and < 0.01 respectively. Optimal model hyperparameters were selected for each chromosome using 5-fold cross-validation to identify the best combination of xgBoost model parameters^104^ for max_depth (5, 10 or 15), min_child_weight (6, 8, or 10), and subsample (0.6, 0.8, 1), with learning_rate = 0.05, min_split_loss = 10, and scale_pos_weight = 1. As recommended, separate models were trained and used for predicting links in each of the six neural cell types analysed.

We evaluated pgBoost predicted SNP-gene links (those scoring in the top 5 % in any cell type) across a range of validation datasets expected to be enriched in true variant-gene pairs, using precision-recall curves given the low prevalence of positive cases^105^. Validation data included: SNP-gene pairs showing significance within cell type-specific eQTL and colocalisation results from independent human brain snRNA-seq studies^22,46^ and SNP-gene pairs comprising credible set lead SNPs and their corresponding target genes prioritised by the Open Targets Locus-to-Gene algorithm^47,48^. Open Targets variant-gene pairs were downloaded from the Open Targets Platform for three GWAS studies: BMI (GCST90691769), AD (GCST90027158) and schizophrenia (GCST90128471) using the European ancestry cohort results where available^48,68,91,106^. Precision-recall curves and reported metrics (Fig. S17, Table S8) were obtained using the R package PRROC^105^, and enrichment calculated as precision / baseline prevalence of true SNP-gene links in each dataset.

## Acknowledgements

The present study was supported by opportunity pool funds from the Accelerating Medicines Partnerships Program in Common Metabolic Diseases (AMP CMD), US National Institutes of Health grant UM1DK105554, and the European Union’s Horizon Europe program (Youth-GEMs, grant agreement no. 101057182). N.E.M. is supported by the MRC (MR/W017156/1). J.M. is supported by the UK Human Functional Genomics Initiative funded by the MRC (MR/Z000068/1). This project utilised equipment funded by the Wellcome Trust (Multi-User Equipment Grant award number 218247/Z/19/Z) and the UK Medical Research Council (MRC) Clinical Research Infrastructure Initiative (award number MR/M008924/1). This study was also supported by the National Institute for Health and Care Research Exeter Biomedical Research Centre. The views expressed are those of the author(s) and not necessarily those of the NIHR or the Department of Health and Social Care.

We acknowledge the Oxford Brain Bank, supported by the Medical Research Council (MRC), Brains for Dementia Research (BDR) (Alzheimer Society and Alzheimer Research UK), Autistica UK and the NIHR Oxford Biomedical Research Centre. We would like to thank the South West Dementia Brain Bank (SWDBB), their donors and donor’s families for providing brain tissue for this study. Tissue for this study was provided with support from the BDR programme, jointly funded by Alzheimer’s Society UK and Alzheimer’s Society. The SWDBB is further supported by BRACE (Bristol Research into Alzheimer’s and Care of the Elderly).

## Consortia

The members of the Youth-GEMs Consortium are Bart Rutten, Sinan Gülöksüz, Therese van Amelsvoort, Lotta-Katrin Pries, Bochao Danae Lin, Angelo Arias-Magnasco, Erika van Hell, Mary Cannon, David Cotter, Melanie Föcking, Subash Raj Susai, Elisabeth Binder, Jim van Os, Jeroen Pasterkamp, Anna Wiersema, Marco Boks, Winni Schalkwijk, Karim Lekadir, Esmeralda Ruiz Pujadas, Noussair Lazrak, Ian Kelleher, Jenni Leppänen, Valentina Kieseppä, Simona Karbouniaris, Lisette van der Poel, Mariël Kanne, Marijke Kolk, Hanske Douwenga, Jordi Rambla, Arcadi Navarro, Liina Nagirnaja, Aldar Cabrelles Munoz, Lauren A. Fromont, Jaanus Harro, Triin Kurrikoff, Reigo Reppo, Dejan Stevanovic, Aleksa Milevic, Jasna Jancic, Marija Nikolic, Maria Bulgheroni, Laura Giani, Margherita La Gamba, Tomislav Franic, Mia Plenkovic, Covadonga M. Diaz-Caneja, Celso Arango, Marta Ferrer-Quintero, Renzo Abregú-Crespo, Emily Guerra-Blacio, Nuria Martín-Martínez, Christel Middeldorp, Enda M. Byrne, Swathi Hassan Gangaraju, Sushma Marla, Jonathan Mill, Eilis Hannon, Emma Dempster, Philippa Wells, Robin Murray, Andrea Danese, Alexander L. Richards, Lucy Riglin, and Michael C. O’Donovan.

## AI

During the preparation of this work, the author(s) used Claude and GitHub Copilot to provide feedback on writing and code. After using this tool/service, the author(s) reviewed and edited the content as needed and take(s) full responsibility for the content of the published article.

