## Supplementary Figures for "A single-nuclei multiomics resource across four brain regions prioritises human neural cell types influencing brain-related traits"

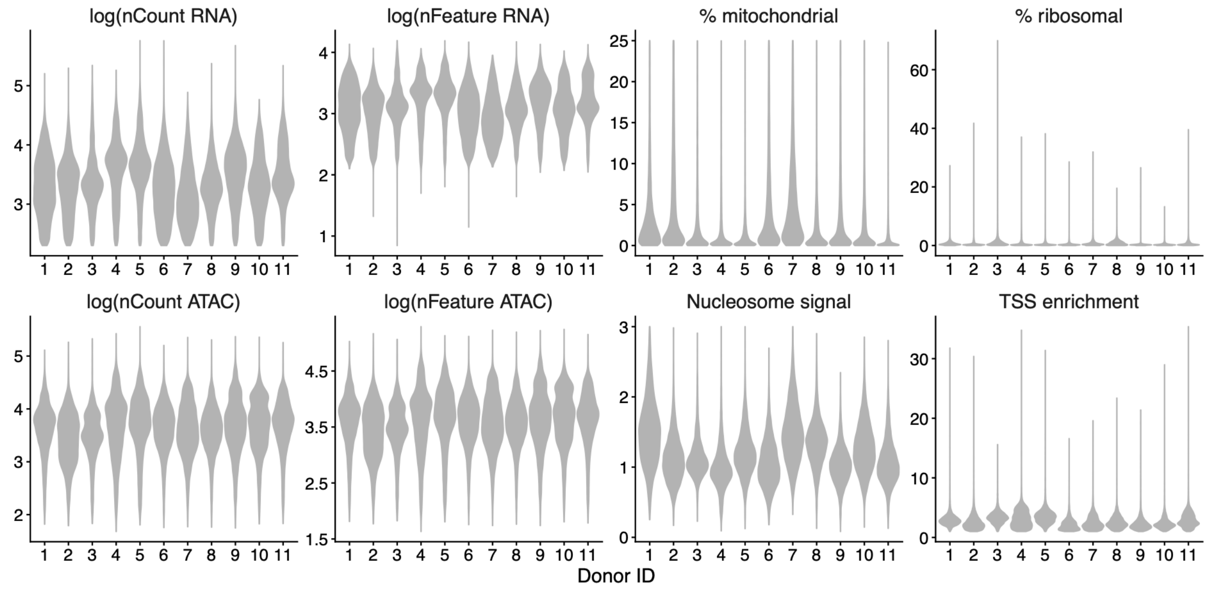


**Figure S1: Quality control metrics for single-nuclei multiome data**. Distribution of values for RNA (*top*) and ATAC (*bottom*) quality control metrics per nuclei across data from each donor. Metrics include: total number of RNA or ATAC counts per nuclei (nCount RNA/ATAC), number of detected genes (nFeature RNA), number of accessible chromatin regions (nFeature ATAC), percentage of reads annotated to mitochondrial or ribosomal reads, nucleosome signal and transcription start site (TSS) enrichment[1].

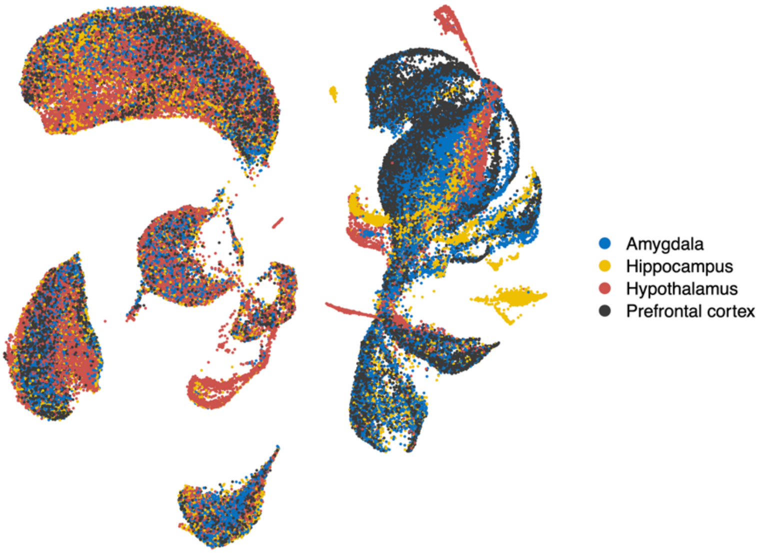


**Figure S2: UMAP visualisation of snRNA-seq data coloured by brain region**. UMAP reduction calculated from RNA data, with each nuclei coloured according to the brain region of origin.

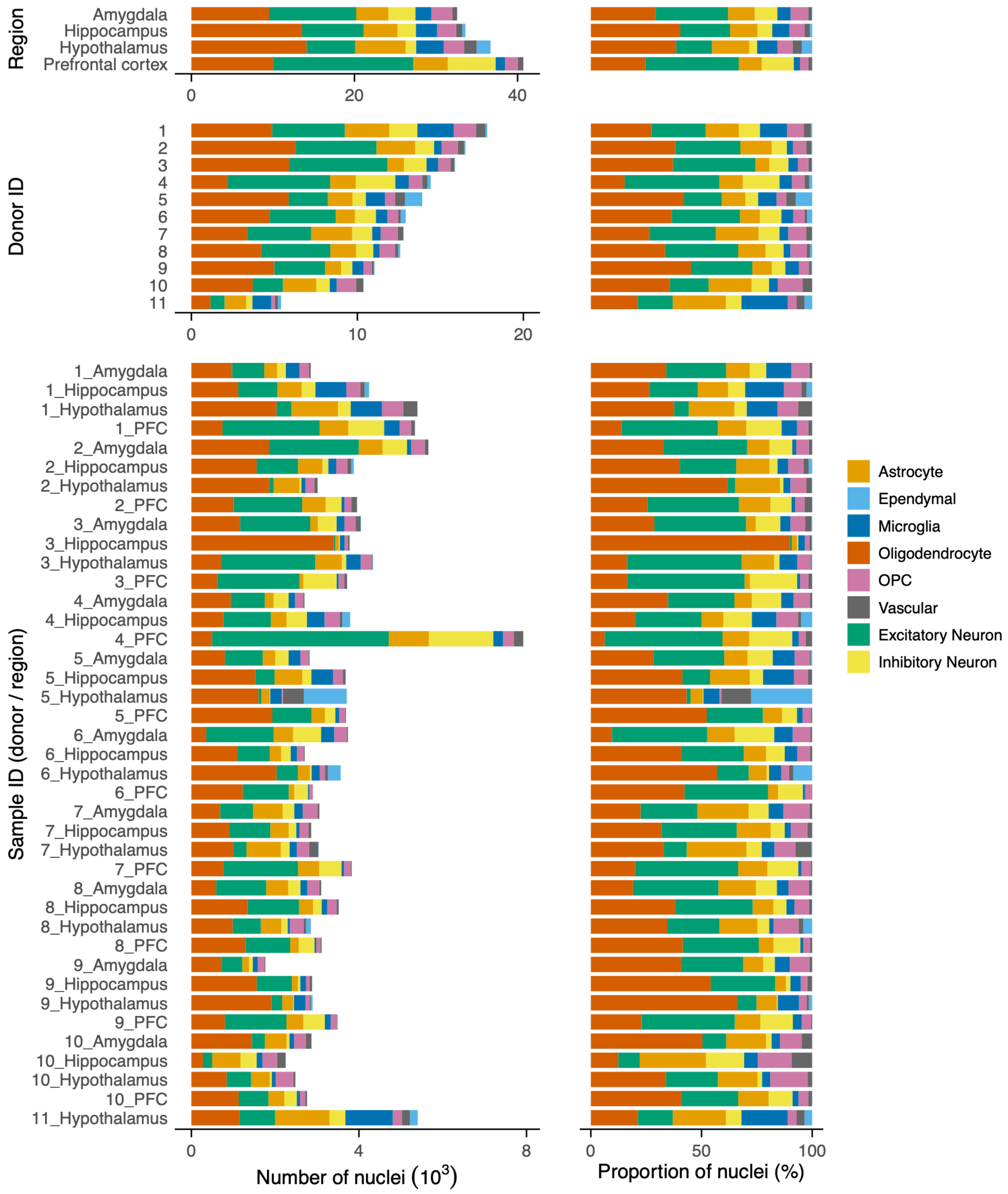


**Figure S3: Cell type composition of samples**. Number (*left*) and proportion (*right*) of total nuclei sampled from each brain region, donor or sample (region / donor combination) that are annotated to each major cell type.

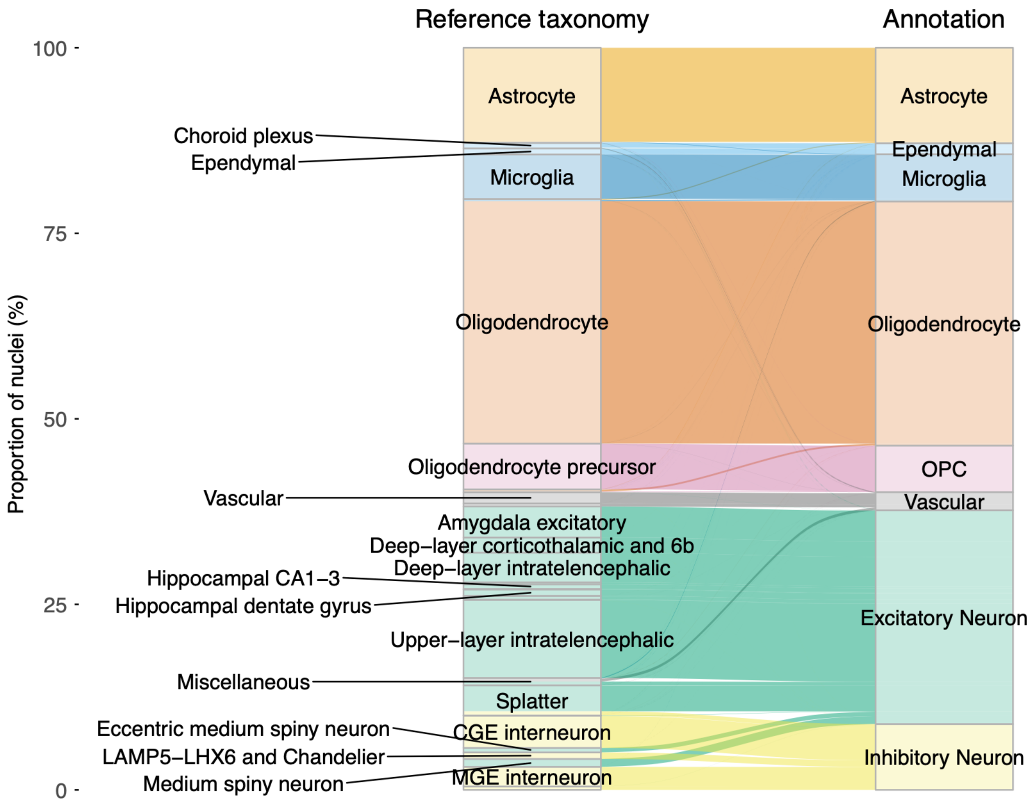


**Figure S4: Comparison of reference-based and manual cell type annotation.** Alluvial plot linking MapMyCells-derived human brain reference taxonomy supercluster labels (*left*) to manually assigned major cell type labels based on marker gene expression (*right*). Bar graphs indicate the proportion of nuclei assigned to each cell type group, with the flow chart illustrating label correspondence between annotation methods.

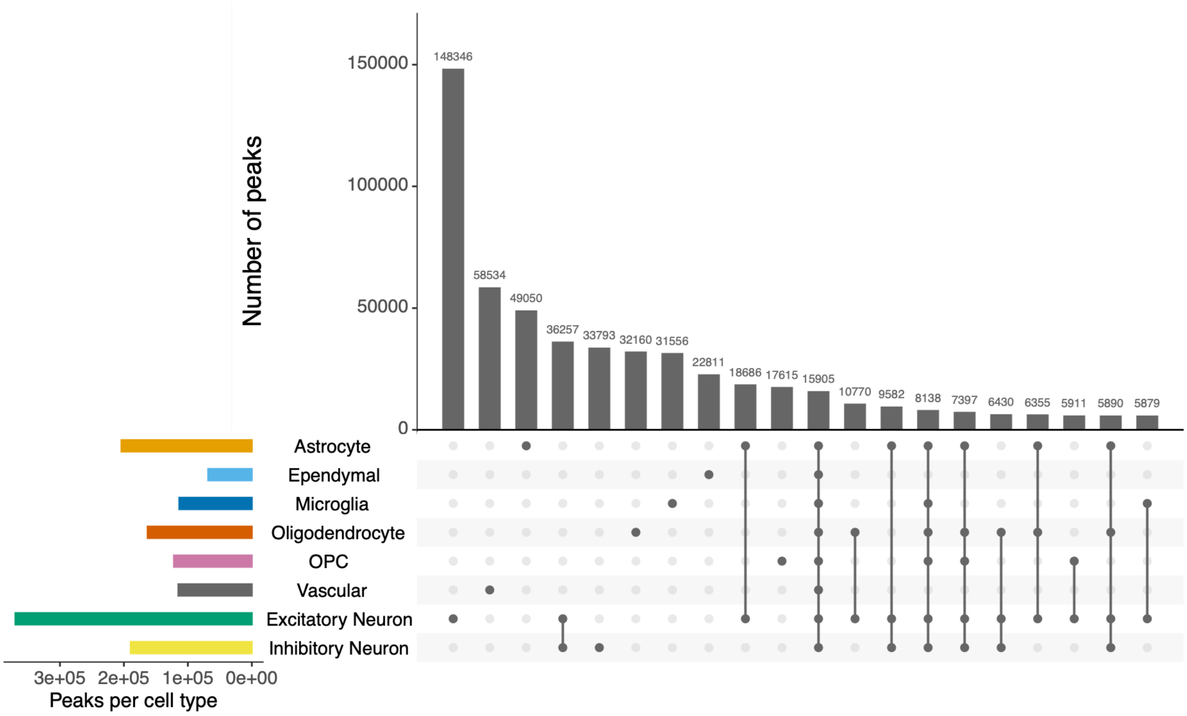


**Figure S5: Cell type-specificity of ATAC peaks**. UpSet[2] plot showing the total number of peaks detected in each cell type (horizontal bars), and the number of peaks (vertical bars) unique to or shared by cell type combinations (indicated by dots below each bar).

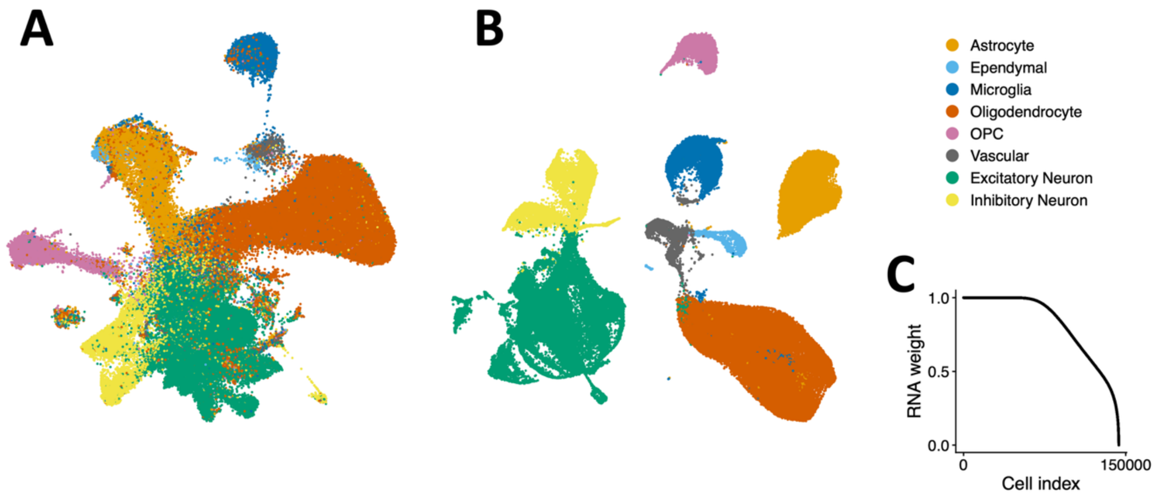


**Figure S6: Weighted-nearest neighbour (WNN) analysis of RNA and ATAC modalities**. UMAP visualisations of single-nuclei data with UMAP reduced dimensions calculated based on (A) ATAC features only or (B) a weighted combination of both RNA and ATAC features; colours indicate the RNA-based major cell type annotations for nuclei. (C) Relative weight of RNA modality for each cell learned by the WNN algorithm[3].

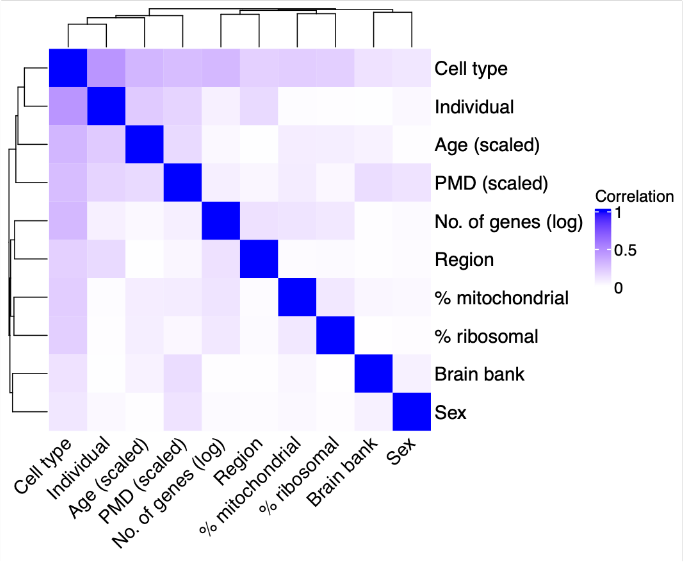


**Figure S7: Correlations between variance partitioning model covariates.** Heatmap indicates pairwise correlations between variables, calculated using Canonical Correlation Analysis to accommodate continuous and categorical variables.

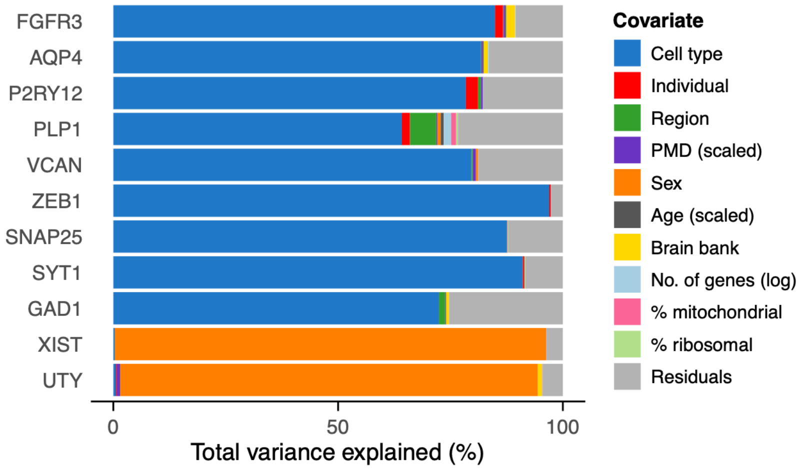


**Figure S8: Proportion of gene expression variance explained by each covariate.** Selected genes represent known neural cell type markers (Table S2) and sex-linked (*XIST*, *UTY*) genes.

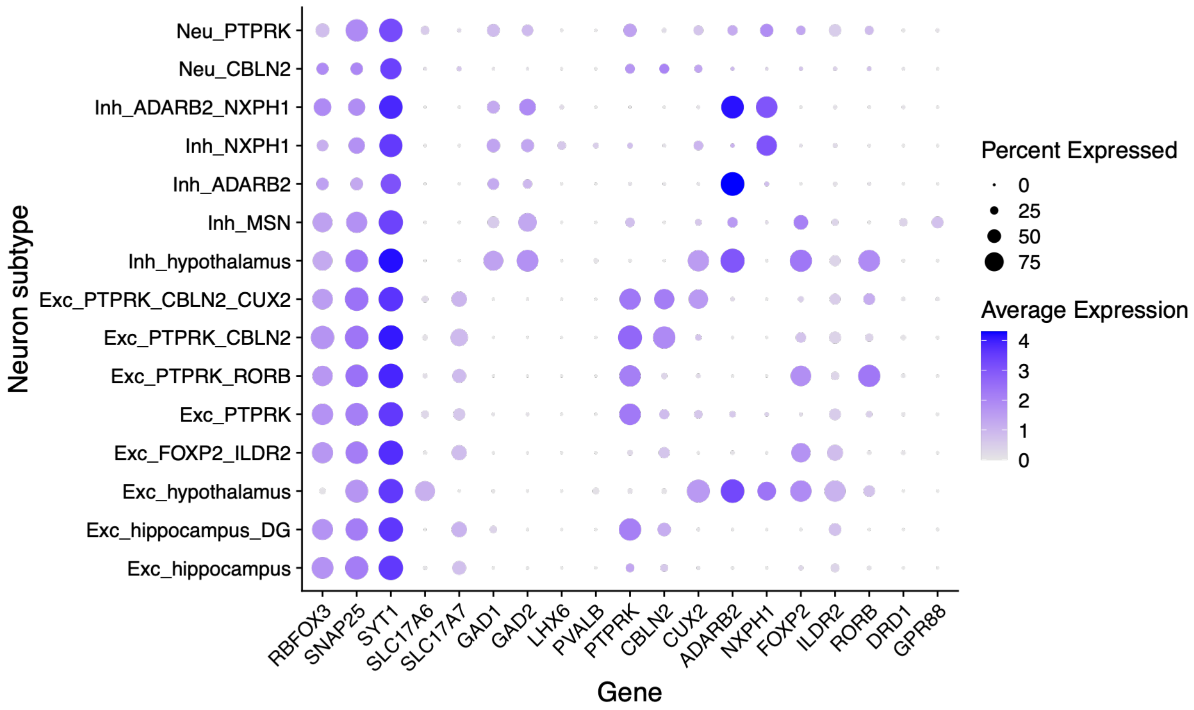


**Figure S9: Expression of selected marker genes in neuron subtypes**. Point size indicates the percentage of nuclei within a given neuron subtype (*row*) which express a given gene (*column*). Point colour indicates the mean (log-transformed) gene expression level across nuclei within that neuron subtype.

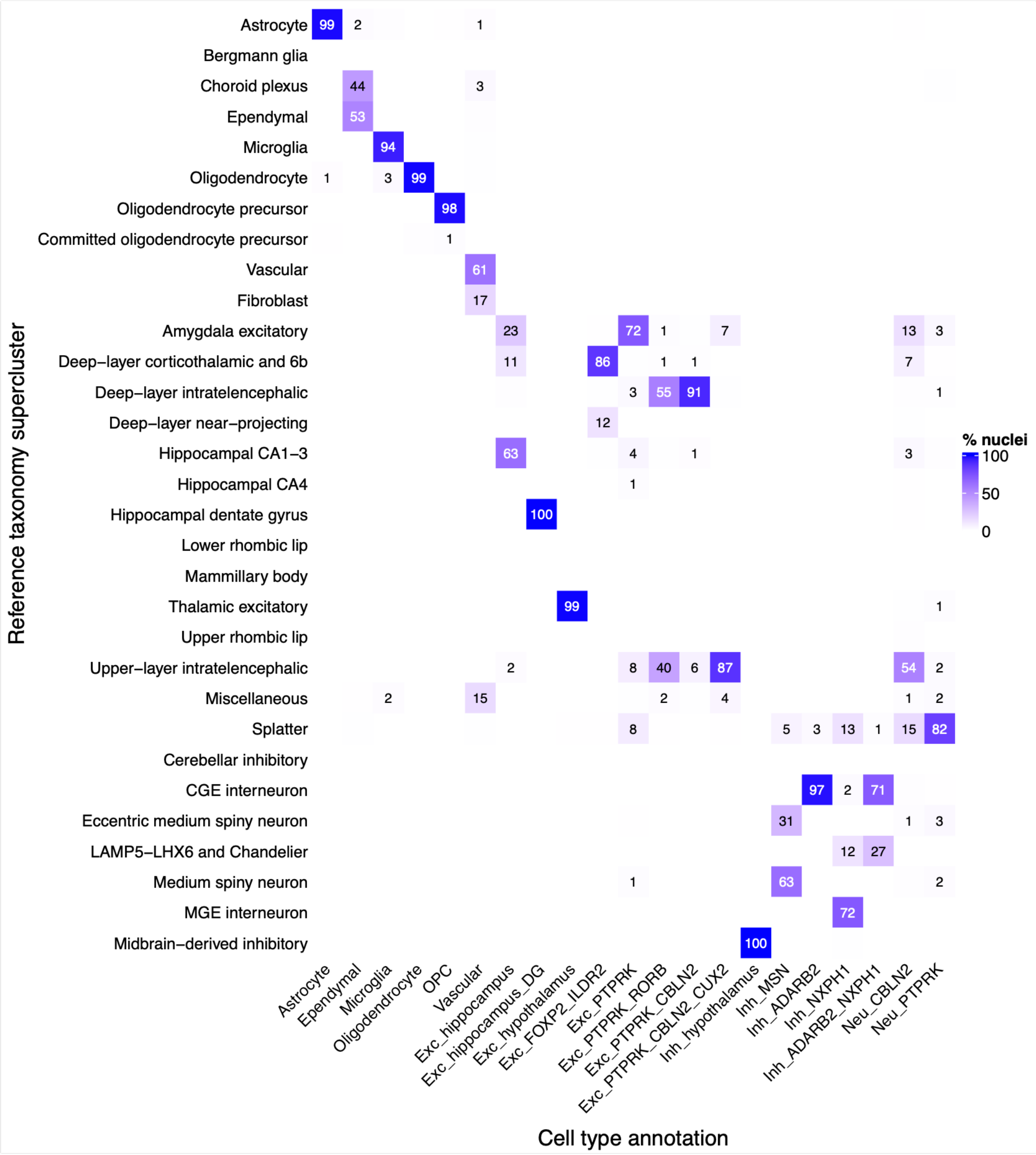


**Figure S10: Comparison of reference-based and manual cell type and neuron subtype annotations.** Heatmap showing the percentage of nuclei within each manually annotated cell type group (*columns*) that are mapped to each reference taxonomy supercluster (*rows*).

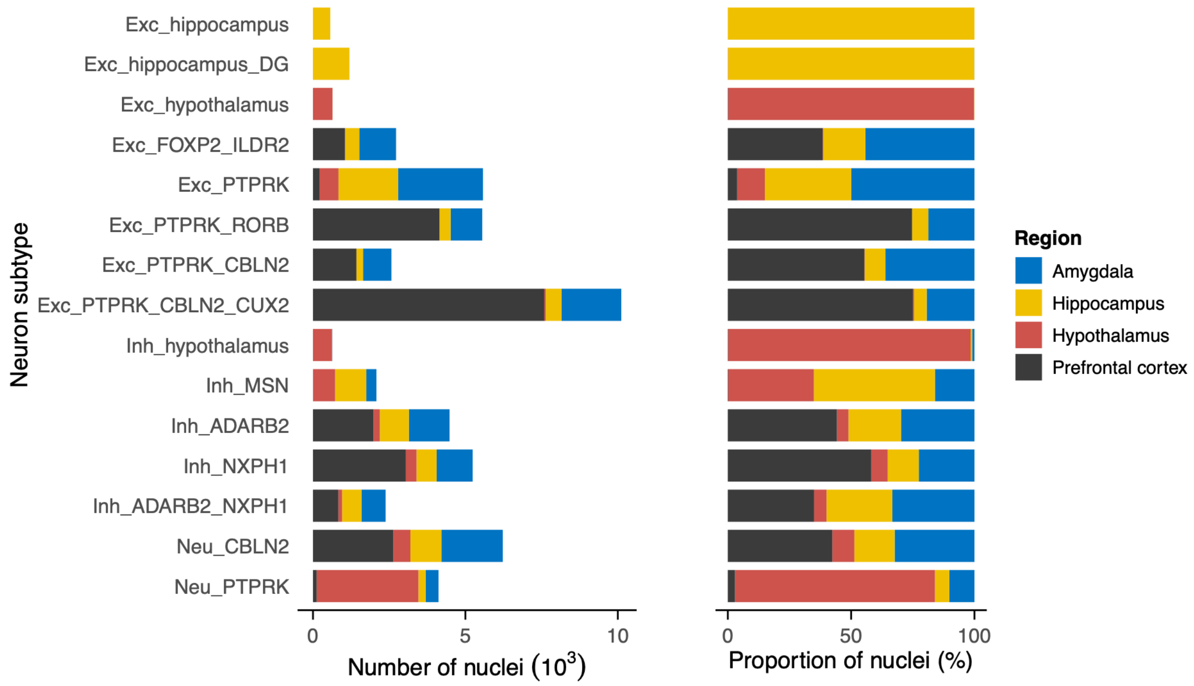


**Figure S11: Neuronal subtype composition of sampled brain regions**. Number (*left*) and proportion (*right*) of total nuclei sampled from each brain region annotated to each neuron subtype.

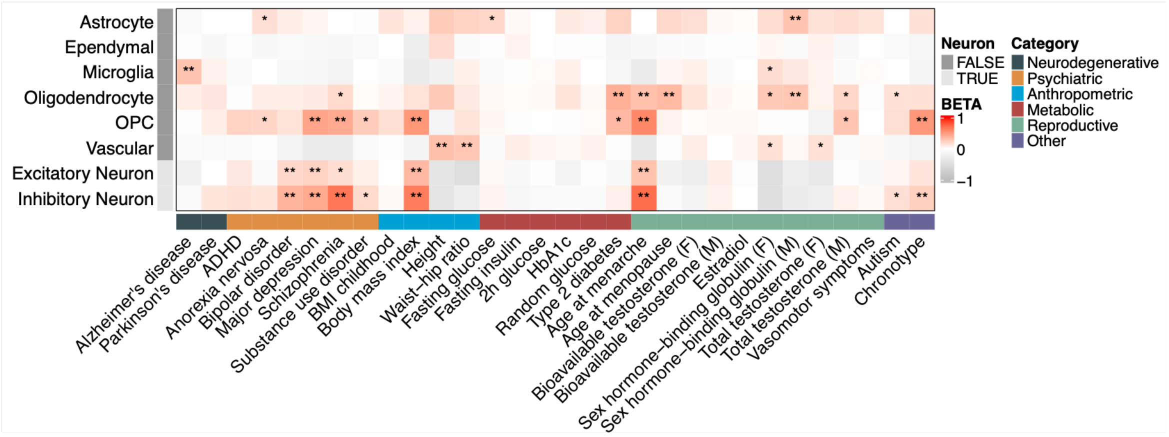


**Figure S12: Association between cell type specific gene expression specificity and brain-related and non-neurological traits**. MAGMA gene property analysis at major cell type level for brain-related and selected metabolic and reproductive traits. Heatmap shows effect sizes (*beta*) for one-sided enrichment tests; * and ** indicate FDR-corrected *p*-values < 0.05 and 0.01, respectively.

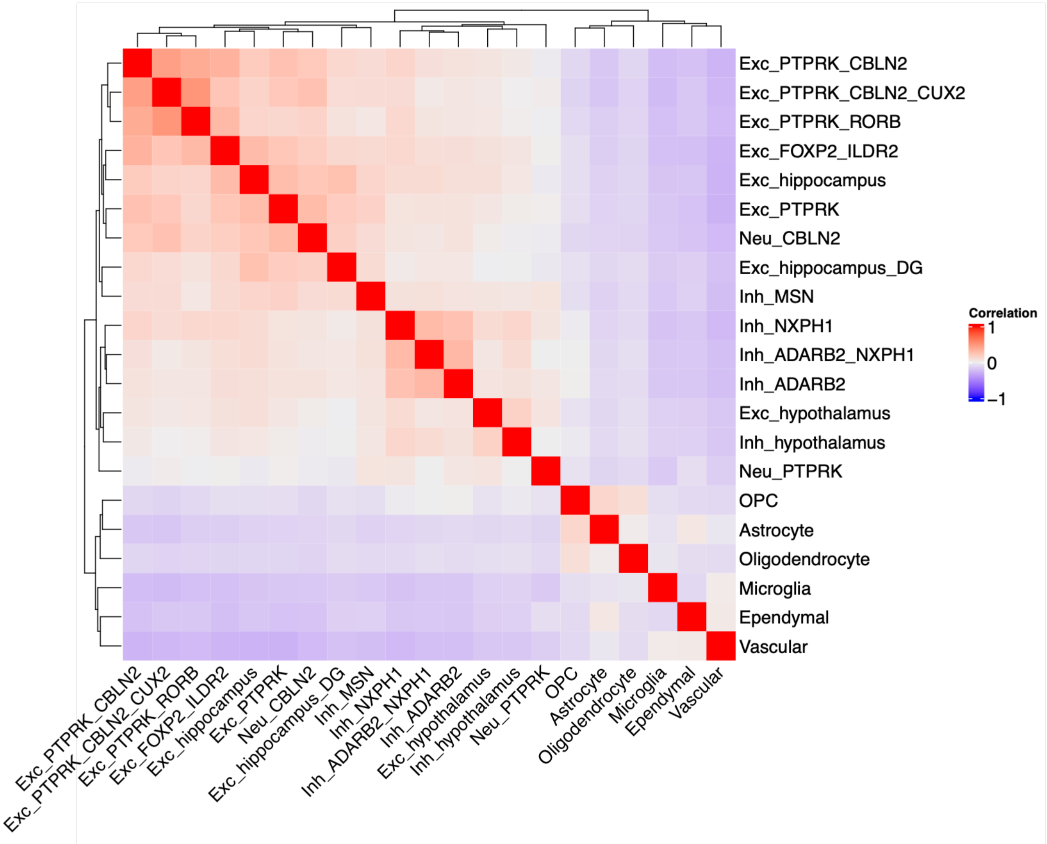


**Figure S13: Correlations of cell type gene specificity profiles**. Pairwise (Pearson) correlation between cell type gene specificity profiles for non-neuronal cell types and neuronal subtypes.

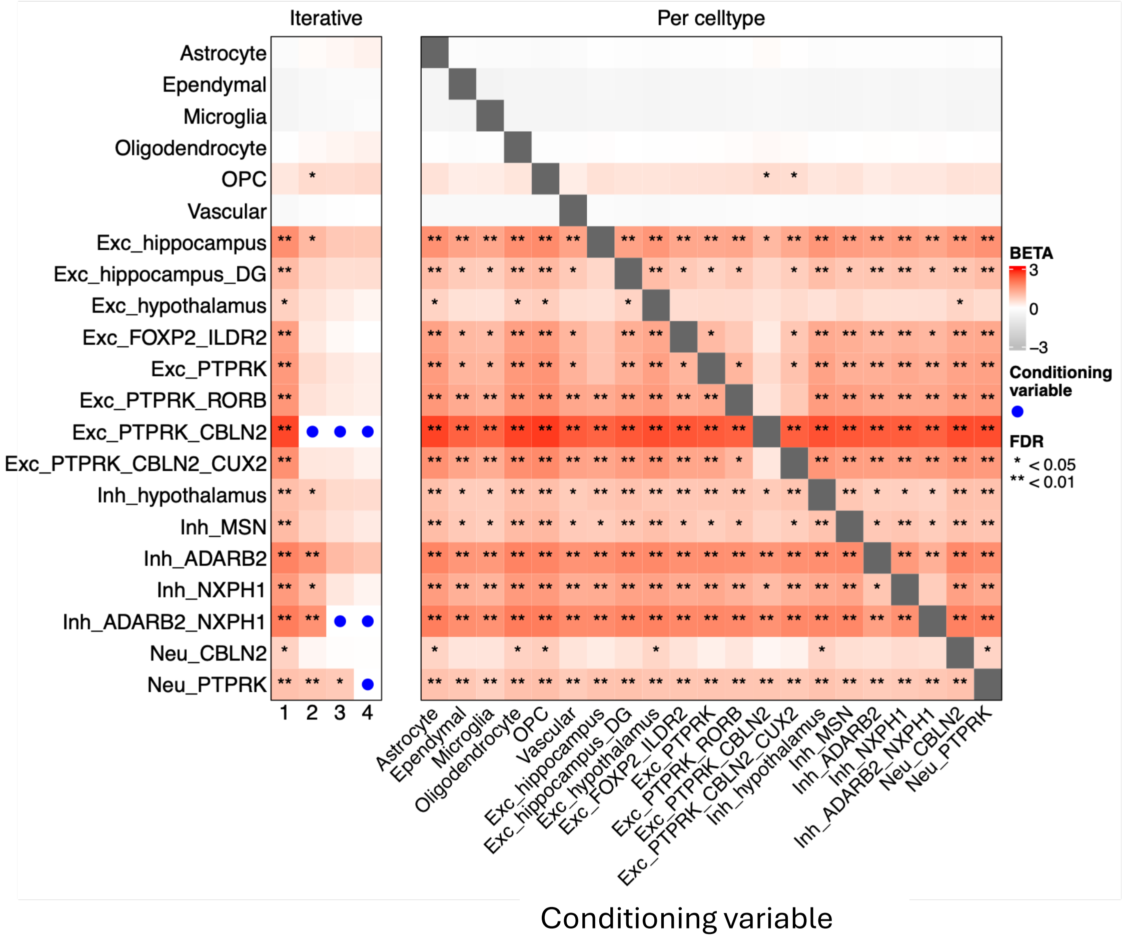


**Figure S14: Conditional MAGMA enrichment analyses for BMI**. MAGMA gene property enrichment analyses for BMI, conditioned on cell type. *Left panel*: iterative conditional analysis results are reproduced from Fig. 3B; in successive rounds, the most significant cell type from the prior round is added to the conditioning set. *Right panel*: per cell type conditional analyses results, where each cell type is individually tested as the conditioning variable.

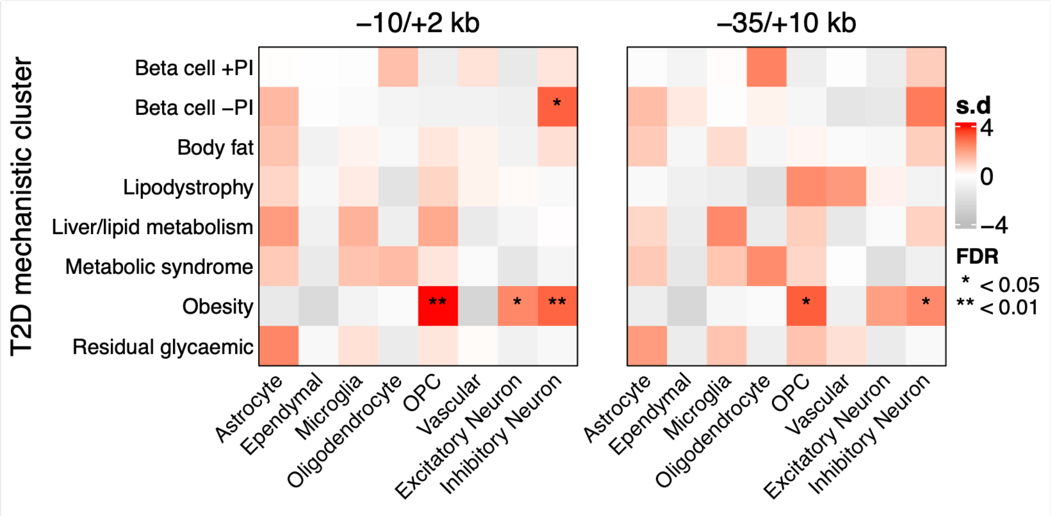


**Figure S15: Expression enrichment of T2D mechanistic cluster-associated gene sets in neural cell types.** Heatmaps show cell type specific expression enrichment of gene sets associated with mechanistic cluster variants from Suzuki et al[4]; gene sets were created by identifying genes where cluster variants overlapped genomic windows of 10 kb upstream / 2 kb downstream (*left panel*) or 35 kb upstream / 10 kb downstream (*right*) around genes (see Methods). Colour indicates standard deviations from the mean during EWCE[5] bootstrapping tests; asterisks indicate cell types showing FDR-significant enriched expression of a given gene set.

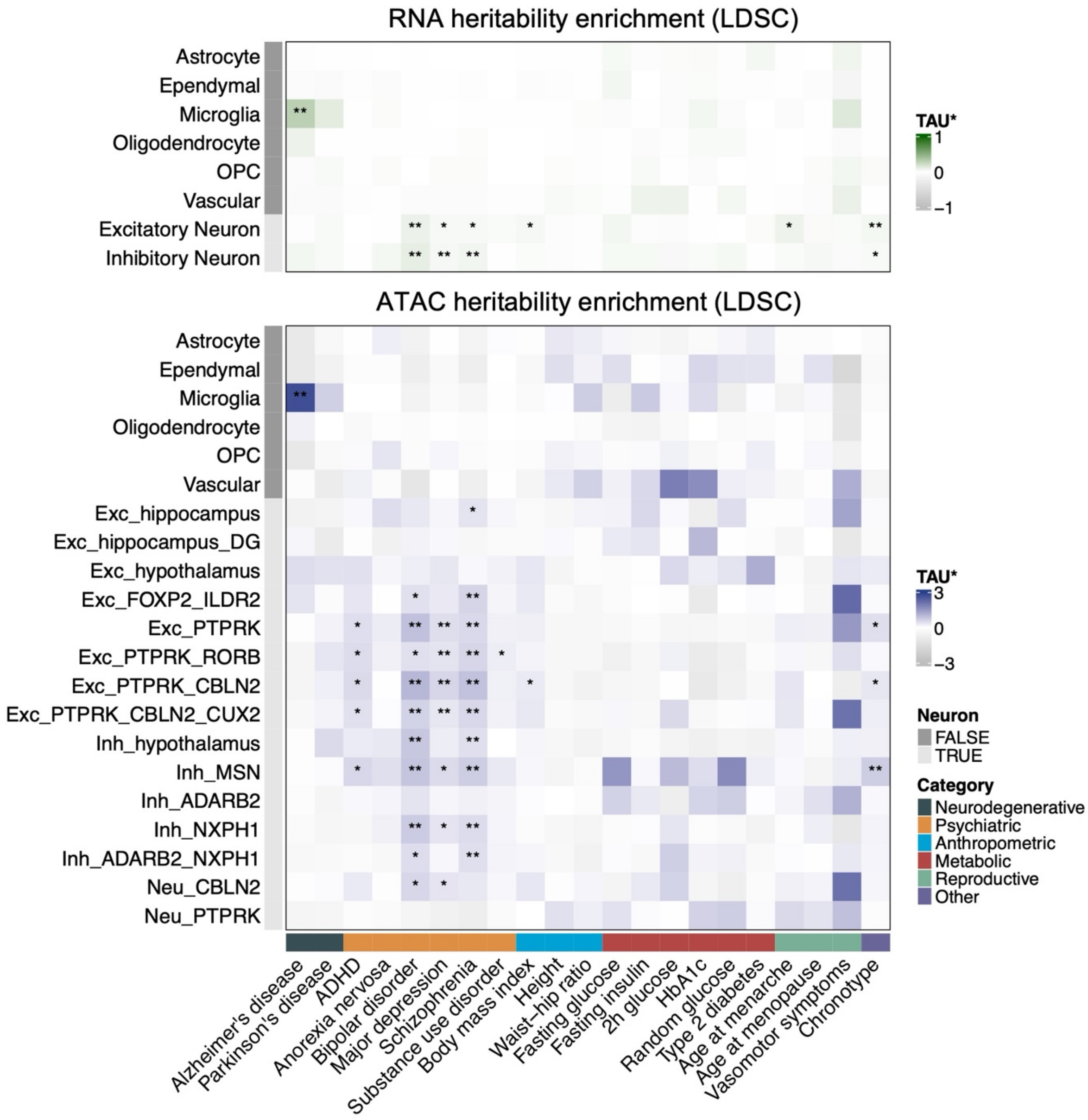


**Fig S16: sLDSC cell type specific heritability enrichments.** Heatmaps show sLDSC heritability enrichments within regions surrounding genes showing cell type specific expression (*upper panel)* or within open chromatin regions showing cell type or neuronal sub type specificity (*lower panel)*. Heatmap shows standardised effect sizes (*tau**); * and ** indicate FDR-corrected one-sided *p*-values < 0.05 and 0.01, respectively, calculated from LDSC coefficient *z*-scores.

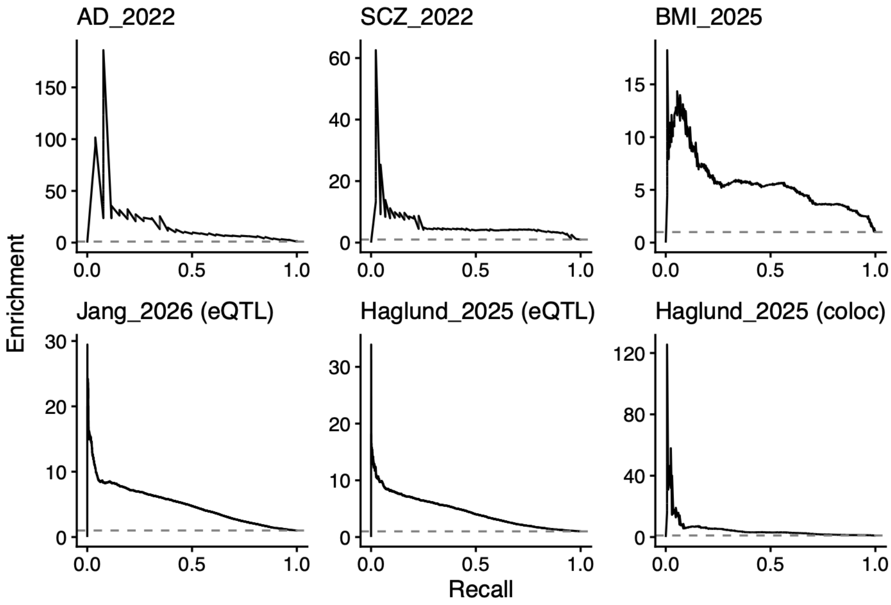


**Figure S17: Enrichment-recall curves to evaluate predicted variant-gene links compared to reference datasets.** Enrichment (precision / baseline prevalence) across recall values for selected validation datasets comprising variant-gene pairs from eQTLs identified using human brain snRNAseq data[6,7], colocalised eQTLs and risk variants for brain-related traits[6], and GWAS credible set lead variants and their prioritised target genes[8]; see Table S8 for details.

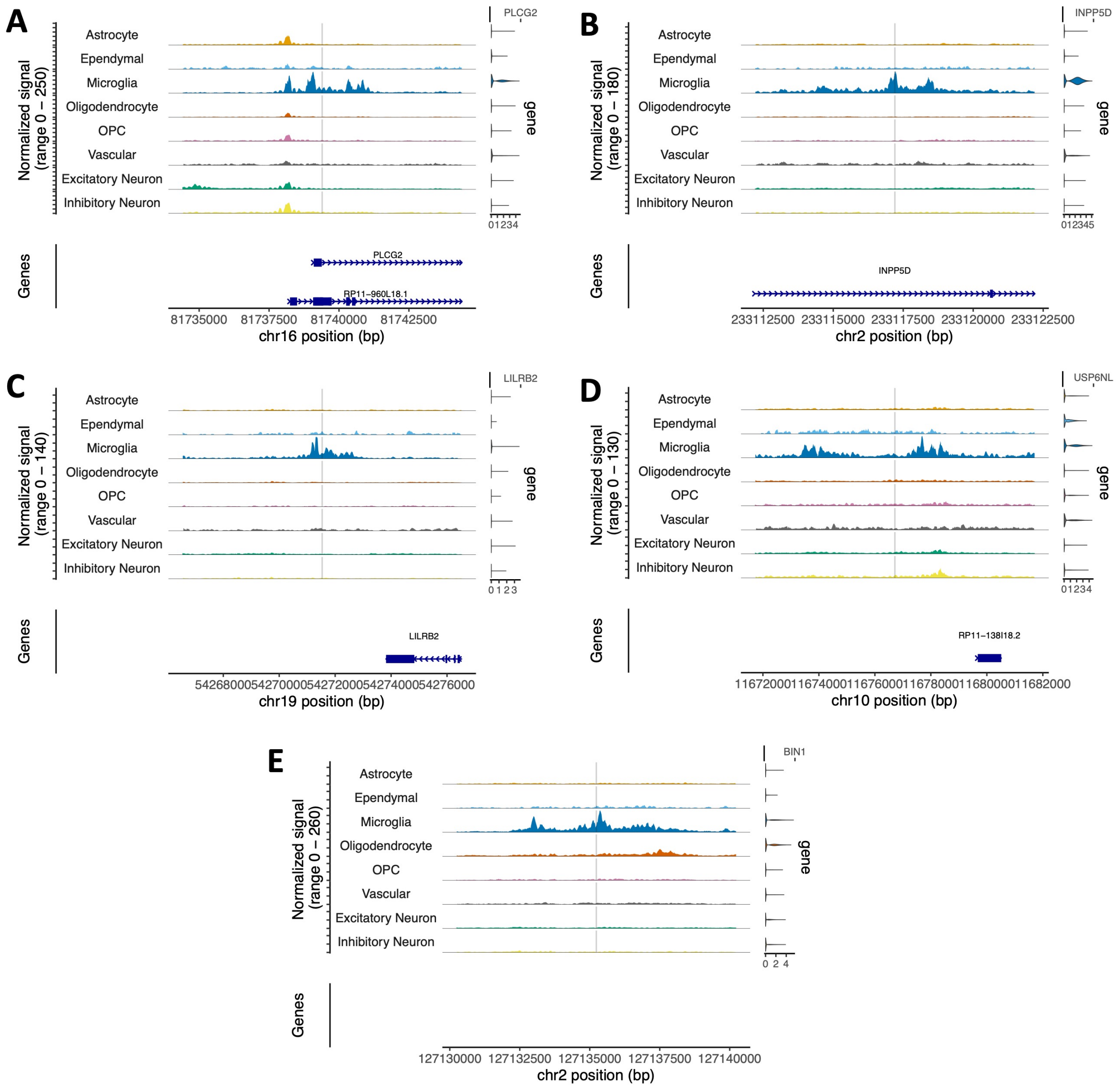


**Figure S18: Linking Alzheimer's disease risk variants to candidate target genes.** (A-E) Additional examples to supplement Fig 4. Coverage plots showing normalised chromatin accessibility signal across major cell types at selected genomic regions. Tracks are centered on regions surrounding genetic variants prioritised by pgBoost, and violin plots (right) show cell-type specific gene expression profiles of the corresponding target genes. In each case the variants represent the lead SNP from an Alzheimer's disease GWAS credible set identified by Open Targets and the same genes are prioritised by Open Targets Locus-to-Gene predictions[8,9]. Further independent evidence supports the relevance of these putative enhancer-gene links: (A) *PLCG2* expression is increased in AD and correlates with amyloid burden[10], and variant rs12446759 (chr16:81739398, hg38) overlaps long non-coding RNA RP11-960L18.1 which lies upstream of canonical *PLCG2* and forms a fusion transcript with *PLCG2*[11]. The AD-risk locus encompassing rs12446759 colocalises with gene expression, transcript usage ad splice junction usage QTLs[11]. (B) *INPP5D* encodes the SHIP1 protein implicated in regulating microlial phagocytosis and immune response and harbours several AD-associated genetic variants[10]. Intronic variant rs10933431 (chr2:233117202) falls within a putative microglia enhancer associated with *INPP5D* via enhancer-promoter mapping analyses[12,13], and at heterozygous loci impacts allelic imbalance in accessible chromatin in iPSC-derived microglia[14]. (C) Supporting functional evidence for variant rs7254645 (chr19:54271535) specifically is lacking, but *LILRB2* is assigned to its credible set by Open Target's Locus-to-Gene model with high confidence (L2G score = 0.893) including support from molecular QTL data[8,9], and LILRB2's microglial amyloid beta receptor function has been implicated in AD[15,16]. However, multiple independent signals have been prioritised in the surrounding region by different GWAS and fine-mapping studies so characterisation of this locus remains uncertain[8,15,17]. (D) rs7912495 (chr10:11676714) is one of multiple variants prioritised by fine-mapping within colocalised AD-risk and microglial *USP6NL* eQTL loci[7,18] and falls in a microglial enhancer associated with the *USP6NL* promoter by proximity-based ligation assays[12]. (E) rs6733839 (chr2:127135234) falls in a microglial enhancer linked to *BIN1* by enhancer-gene mapping and has been shown to affect chromatin accessibility and *BIN1* expression in molecular QTL studies[7,11,12,14,19,20]. This variant is predicted to create a MEF2C transcription factor binding site[21], with iPSC studies showing enhancer deletion decreases *BIN1* expression[12].
